# 300 billion years of angiosperm evolution at risk of extinction

**DOI:** 10.1101/2025.04.29.651190

**Authors:** Félix Forest, Ruth Brown, Sven Buerki, Jonathan F. Colville, Justin Moat, Eimear Nic Lughadha, Nisha R. Owen, Domitilla C. Raimondo, Malin Rivers, James Rosindell, Barnaby E. Walker, Steven P. Bachman, Sebastian Pipins, Rikki Gumbs, Matilda J.M. Brown

## Abstract

Extinction results in not only loss of species, but also loss of the unique evolutionary history that they represent and the irreplaceable features they exhibit. There is broad consensus regarding the necessity to optimise the preservation of the tree of life by including evolutionary information in conservation prioritisation, a notion also endorsed by major policy frameworks^1–4^. However, evolutionarily-informed prioritisations are lacking for most plants, resulting in a taxonomic imbalance in the evolutionary information incorporated in global biodiversity analyses, which has undermined conservation for decades. Here, we use comprehensive species-level phylogenetic trees, and extinction risk estimates, to generate the first global assessment of angiosperm evolutionary history at risk, and to identify phylogenetically-informed conservation priorities for the world’s flowering plants. We estimate that more than one fifth of angiosperm evolutionary history is at risk of extinction in the short term. Using the Evolutionarily Distinct and Globally Endangered^5,6^ approach, we identify 9,945 threatened plant species that disproportionately account for total evolutionary history at risk. Species and area prioritisations incorporating evolutionary history are urgently needed to correct imbalances between plants and animals, monitor the effectiveness of conservation efforts, and optimise conservation resource allocation in the face of increasing human pressures on Earth’s biodiversity.

## Main

Not all species are equal in evolutionary terms^7^. A species isolated on the tree of life (e.g. *Amborella trichopoda*, the sole extant representative of the order Amborellales), embodies more unique evolutionary history than a recently diverged species with several close relatives (e.g. one of >3,000 *Astragalus* spp). Thus, the successful preservation of biodiversity would include maximising the protection of the evolutionary lineages that led to the diversity of life on Earth today. Optimising the preservation of evolutionary history should therefore form an integral part of an effective and future-proof conservation agenda^8^. The maintenance of these evolutionary lineages is expected to also conserve substantial feature diversity, i.e. the variety of traits of species from which current and future benefits for society and nature stem^9–13^. In fact, given the paucity of trait information for large groups such as flowering plants (i.e. angiosperms), accounting for evolutionary history may be the only practicable approach for the evaluation of feature diversity in global analyses. The effective conservation of multiple facets of biodiversity depends on the continued existence of the diverse evolutionary lineages of all life on Earth. While the necessity to optimise the preservation of the tree of life is widely advocated and generally accepted, both in the scientific literature and in global policy^3,4^, in practice it often remains overlooked^14^.

Various prioritisation approaches have been proposed that attempt to capture the evolutionary dimensions of biodiversity. Trends in both the overall expected loss of evolutionary history, measured as the length of all the branches linking a set of taxa on a phylogenetic tree (i.e., phylogenetic diversity, PD^15^), and the conservation status of the most evolutionarily distinct species are now recognised in global biodiversity policy initiatives^1,2^. The Evolutionarily Distinct and Globally Endangered (EDGE) approach^6^, which ranks species according to how much evolutionary history would be safeguarded should a species be conserved, has been used to direct conservation resources for several animal groups^6,16–19^ and gymnosperms^20,21^. Despite its application to some groups of angiosperms (e.g. *Erica*^22^), EDGE has not yet been applied to flowering plants at a global scale. The application of the EDGE approach to the angiosperms as a whole will offer critical information require for the reporting of the two tree of life indicators recognized by the Kunming-Montreal Global Biodiversity Framework (GBF)^1^ of the Convention on Biological Diversity, the ‘Phylogenetic Diversity Indicator’ to monitor the global status of the tree of life, and the E‘DGE index’ to assess the conservation of evolutionary unique species^3^.

Approximately 335,000 species of angiosperms are known to science, forming the living dominant components of virtually all terrestrial ecosystems^23^ and underpinning the survival of countless other species of plants, fungi, animals, and microorganisms. Machine learning predictions from models trained on IUCN Red List assessments indicate that more than 2 in 5 plants are threatened with extinction^24^, but only ∼20% of plant species have global IUCN Red List assessments^25^, compared to >80% in all major vertebrate groups, hindering the inclusion of plants in most conservation prioritisation approaches. Fortunately, it is now possible to produce EDGE rankings for groups with incomplete data, as uncertainty in both phylogeny and extinction risk are handled in the updated EDGE2 protocol^5^. Recent developments in species imputation methods^26,27^, coupled with comprehensive and curated lists for described species (e.g. World Checklist of Vascular Plants^28^), enable the creation of complete species-level phylogenetic trees from incomplete molecular data, especially pertinent to large, relatively data-poor taxonomic groups like angiosperms. EDGE scores are calculated across large sets of such trees, representing a distribution of possible phylogenetic hypotheses. The probability of extinction for each species is assigned by sampling from a distribution generated by fitting a quartic curve through the median values associated with IUCN Red List categories^5^, enabling the inclusion of species lacking published IUCN Red List data-sufficient assessments by sampling uniformly from this curve. The use of extinction risk predictions for species lacking published IUCN Red List assessments allows a more targeted sampling of the distribution of extinction risk values for unassessed and data deficient species.

Here, we produce the first comprehensive, global analysis of conservation priorities for angiosperms based on phylogenetic information and estimate how much their evolutionary history is at risk of extinction. To achieve this, we use a backbone phylogenetic tree based on DNA data and species imputation methods to reconstruct a set of 200 complete species-level phylogenetic trees for 335,497 recorded species of angiosperms. These trees are an order of magnitude larger than the next-largest such phylogenetic tree used for EDGE analyses (i.e. ray-finned fish, Actinopterygii, ∼33,000 spp^29^) and orders of magnitude greater than the largest group of plants for which EDGE scores have been compiled, i.e. gymnosperms (∼1,100 spp^20^). We combine our novel trees with extinction risk estimations for all species based on IUCN Red List categories and Bayesian mixed modelling for species yet to be assessed for the global IUCN Red List. In doing so, we produce a list of priority threatened species representing a substantial amount of evolutionary history at risk (i.e. EDGE List^5^). To illustrate the immediate relevance of such a loss on nature’s contribution to people calculate how much threatened evolutionary history is represented by plants used by humans. Finally, we explore different scenarios of increased extinction risks and how these would affect the amount of evolutionary history at risk and the number of species included in EDGE Lists.

### Global status of evolutionary history

Large-scale estimates of diversity loss are currently hindered by lack of extinction risk information for most species^30–32^. We generated comprehensive estimates of extinction risk for angiosperms, based on geographic distribution, lifeform, and documented human use, to improve the calculation of expected loss of evolutionary history across the entire clade (see Methods). Angiosperms comprise 1,445 billion years (byr) of evolutionary history. More than a fifth of this is at risk (307 byr, 21.2%; median from 200 trees; Fig. 1a) as estimated by complementing data from global IUCN Red List assessments with our extinction risk predictions. This estimate is lower than the 26.7% (386 byr) calculated to be at risk when we omit our extinction risk estimates, treating Data Deficient (DD) and Not Evaluated (NE) species as in Gumbs et al.^5^. This discrepancy is due to differences between the more animal-focussed approach to DD/NE species of Gumbs et al.^5^, in which sampled extinction probabilities of DD/NE species are, on average, equivalent to Vulnerable (VU), given the association between data deficient animals and predicted elevated extinction risks^33^. On the other hand, our extinction risk predictions have sampled extinction probabilities for DD/NE species equivalent to

**Figure 1.**
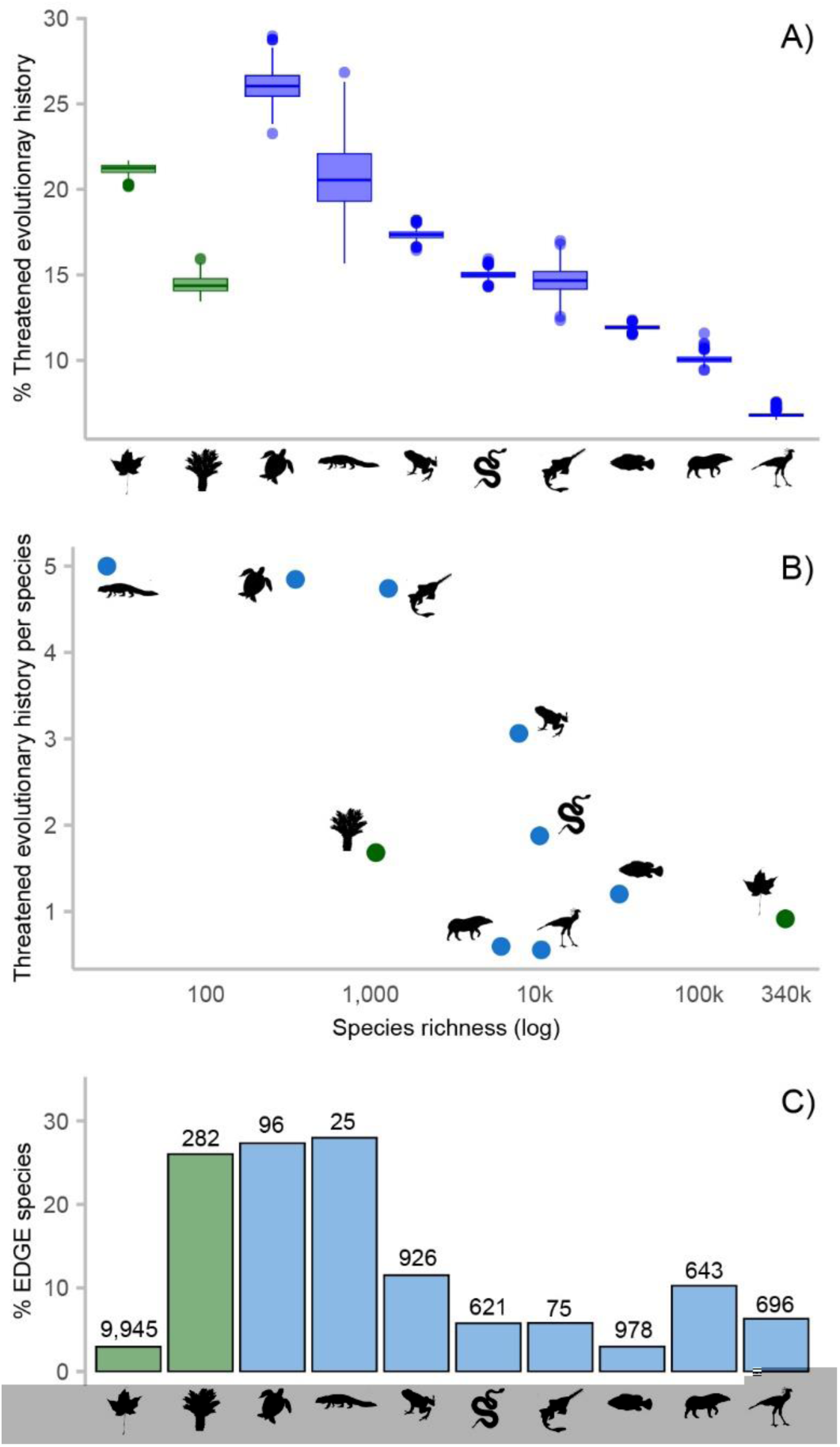
Comparison of threatened evolutionary history and EDGE species numbers across selected groups of plants and animals. A) Proportion (in %) of evolutionary history at risk of extinction in all major groups of plants (green shading) and animals (blue shading) for which this information is available (data from this study, Gumbs et al^19^, and https://www.edgeofexistence.org/), calculated across all replicates (angiosperms, N = 200; gymnosperms, N = 100; vertebrate clades, N = 1000); centre line in boxplots show the median. From left to right: plant groups (angiosperms, gymnosperms) and jawed vertebrate groups (testudines, crocodylians, amphibians, lepidosaurs, chondrichthyans, ray-finned fish, mammals and birds). B) The amount of threatened evolutionary history per species for each group against their total species richness (log scale). C) Proportion (in %) of EDGE species in all major groups of plants and animals for which this information is available (data as in A). Numbers above bars are the number of EDGE species in each group. Groups ordered as in panel A.

Near Threatened (NT) for species determined to be non-threatened and slightly more threatened than VU for threatened species. Given that we predict 57.3% of DD/NE species (163,720 out of 285,697 species) as not threatened, a proportion in line with previous estimates^24^, ignoring extinction risk predictions and relying only on the limited number of published global Red List assessment data results in an inflation of the evolutionary history expected to be lost. These results highlight the value of extinction risk predictions, particularly for large clades with low data coverage like angiosperms. Hereafter, we lead with our results based on the analyses which include extinction risk predictions, commenting on the results obtained using only data-sufficient IUCN Red List assessments where the differences between the two are of particular interest.

The families with the largest proportion of evolutionary history at risk include the Magnoliaceae (34.4%), Pandanaceae (31.7%) and Ancistr ocladaceae (30.6%), among families comprising ≥10 species. The amount of threatened evolutionary history per family is generally correlated with total evolutionary history per family (Fig. S1). The five angiosperm families with the largest amount of evolutionary history at risk are Orchidaceae (26.4 byr; 30,079 spp), Araceae (23.2 byr, 4,363 spp), Rosaceae (17.2 byr, 5,176 spp), Fabaceae (16.7 byr, 22,293 spp) and Myrtaceae (13.8 byr, 6,207 spp). Orchidaceae and Fabaceae are the second and third largest families in terms of species number in angiosperms (after Asteraceae), respectively, and are unsurprisingly among the largest contributors to the total evolutionary history at risk in angiosperms, but the other three families are comparatively small. Collectively, these five families account for almost a third of all angiosperm evolutionary history at risk (31.9%) while comprising about a fifth of total angiosperm species richness (20.3%). The 21.2% of overall angiosperm evolutionary history at risk is substantially greater than the 13% (102 of 782 byr) estimated for jawed vertebrates, ranging from 7% for birds to 26% for turtles and tortoises (Fig. 1a)^19^. Conversely, the average amount of evolutionary history at risk of extinction per angiosperm species (i.e. 0.92 myr) is greater than that reported for mammals and birds, but lower than that of any other clade of jawed vertebrates (Fig. 1b)^19^.

With only one assessment time point we cannot calculate trends in angiosperm extinction risk and associated indicators such as the EDGE index^3^. However, with ongoing increase in extinction risk across major groups^34^, we can expect similar patterns of decline for angiosperms, though the observed decline is rarely as severe as if every species was reassessed and assigned to the adjacent more threatened category. To estimate the impact of continuing declines across angiosperms, we simulated incremental increases in extinction risk for all species under three scenarios, i.e. increase of +0.2, +0.5 and +1 IUCN Red List categories (see Methods and Fig. S2). When recalculating the expected loss of angiosperm evolutionary history under these three scenarios, the proportion of evolutionary history at risk of extinction increased from 21.2% under current extinction risk to 28.0% in the most cautious scenario (+0.2), and 34.9% in the most extreme (+1), exceeding 500 billion years of threatened evolutionary history (details for all scenarios in Table S1). Even the most cautious scenario would result in an additional 40 billion years of threatened evolutionary history compared to the current level. Given that threat is not random on the tree of life^35,36^, some groups will be more affected than others by a global (most likely uneven) increase in probability of extinction, which would probably result in even greater loss of evolutionary history.

### EDGE species

Certain species are responsible for disproportionately large amounts of threatened evolutionary history. Consequently, the conservation of these species, known as EDGE species, can effectively safeguard imperilled branches of the tree of life. Originally, EDGE species were identified as those species that both have above-median EDGE scores (with ≥ 95% confidence) and have been assessed as threatened on the IUCN Red List^5,25^. Under this definition, species assessed as DD or NE cannot be considered EDGE species, irrespective of their EDGE score. However, such a characterisation of EDGE species would overlook many priority species of angiosperm, since more than 80% of species are DD/NE on the IUCN Red List (285,697 out of 335,497 species^37^; but see ‘Alternative lists’ section in Supplementary Material). Ther efor e, alongside the set of ‘strict’ EDGE species based on threatened IUCN Red List categories (i.e. CR, EN, VU, and EW), we also highlight ‘candidate’ EDGE species. These are DD/NE species estimated as threatened in at least 50% of the Bayesian mixed modelling draws (100 out of 200; see Methods) and which are above median EDGE with 95% confidence. If assessed, these species are expected to qualify as EDGE species, but uncertainty in their extinction risk precludes their designation as EDGE species in the strictest sense. Here, we identify 5,869 strict EDGE species and an additional 4,076 candidate EDGE species, for a total of 9,945 EDGE species (Fig. 2). The documentation of candidate EDGE species demonstrates the importance of refining extinction risk probabilities using machine learning approaches, especially in groups where full extinction risk assessment data is largely unavailable and where addressing data gaps would be significantly more costly than applying predictive models^38^. It also highlights a set of species which should be prioritised for a full IUCN Red List assessment.

**Figure 2.**
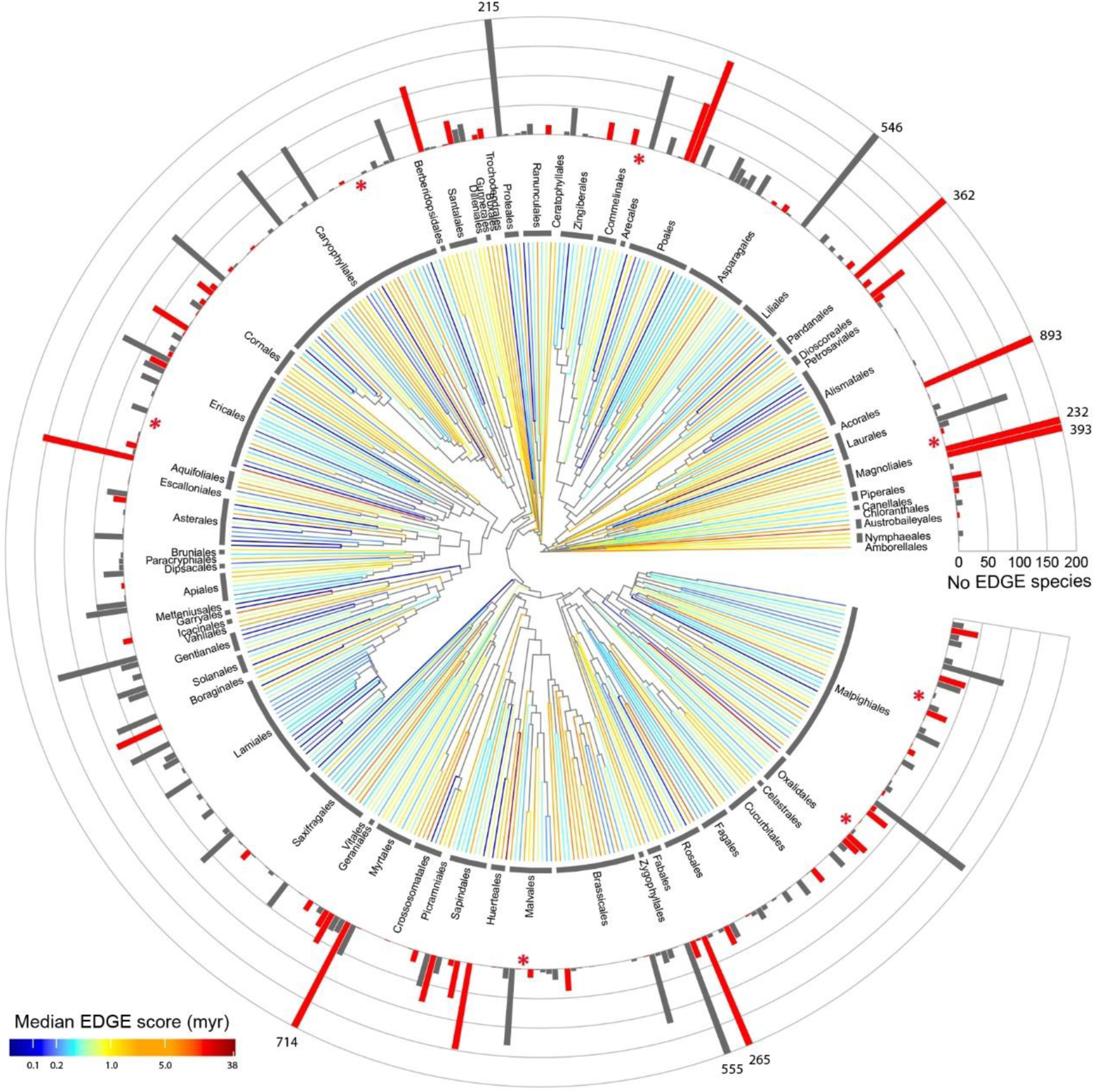
Distribution of EDGE species and EDGE families across the tree of life of angiosperms. Based on one of the 200 species-level trees produced for the present study and pruned to a single representative per family (all 200 trees are identical at the family level); orders only are indicated for clarity. Bars indicate the number of EDGE species in each family (red bars designate EDGE families); EDGE species number are indicated above truncated bars for families with more than 200 EDGE species; red asterisks (*) identify the seven EDGE families with no EDGE species. Terminal branches are coloured according to the median EDGE family score (i.e. mean EDGE score of all species within a family) across 200 trees.

The number of EDGE species of angiosperms is large compared to other clades of plants and animals for which EDGE scores have been compiled (10.2x the number of EDGE ray-finned fish, the next-largest set with 978 EDGE species; Fig. 1c). However, it represents only 2.96% of angiosperm species, a proportion much lower than observed for most other groups, except maybe for ray-finned fish (2.99%; Fig. 1c). On the other hand, the number of species found on the EDGE list represent 16.6% (median obtained from 200 extinction probability weighted trees) of all angiosperm threatened evolution history, which further highlights how EDGE species are disproportionately important for the preservation of the tree of life. Notably, the cumulative sum of EDGE scores for the top 5.9% species with the highest-ranking EDGE scores (median of 19,715 species across 200 compilations) captures 50% of angiosperm threatened evolution history, while the top 9.6% species (6,760 spp) are required to achieve the same coverage for jawed vertebrates^19^. These statistics provide grounds for optimism that directed conservation actions can meaningfully advance the Global Biodiversity Framework targets^1,3^.

Twenty-three of the 25 highest-ranked EDGE angiosperms are all assessed as threatened on the IUCN Red List, the remaining two being estimated to be threatened by our extinction risk predictions. The top EDGE species is *Hondurodendron urceolatum* (Olacaceae) which is categorised as CR on the IUCN Red List; Fig. 3). This dioecious tree is the only species of its genus and is found in scattered populations within a single mountain range in the Parque Nacional El Cusuco in Honduras^39^. While two other species of Olacaceae are ranked 17th and 18th (two species of *Octoknema*), the top 25 EDGE species of angiosperms is dominated by Araceae (seven species, including five of *Amorphophallus*) and Magnoliaceae (six species of *Magnolia*; Fig. 3). The presence of several *Magnolia* and *Amorphophallus* species among the high-ranking EDGE species is most likely due to the combined effect of these being relatively old lineages^40^ which also have a high proportion of threatened species (67.5% and 66.1%, respectively; for *Amorphophallus* this is almost entirely based on extinction risk predictions). Other notable EDGE species in the top 25 include: *Medusagyne oppositifolia* (Ochnaceae; ranked 3rd; CR), a tree found only on the island of Mahé in the Seychelles and known locally as the jellyfish tree^41^; *Afrothismia gesnerioides* (Burmanniaceae; ranked 6th; CR) is a non-photosynthetic mycoheterotrophic herb from the Nyangong forest of Cameroon^42^; and *Gomortega keule* (Gomortegaceae; ranked 11th; EN), the only species of its family, found in a small area on the coast of central Chile, with an edible fruit and wood used as timber^43^.

**Figure 3.**
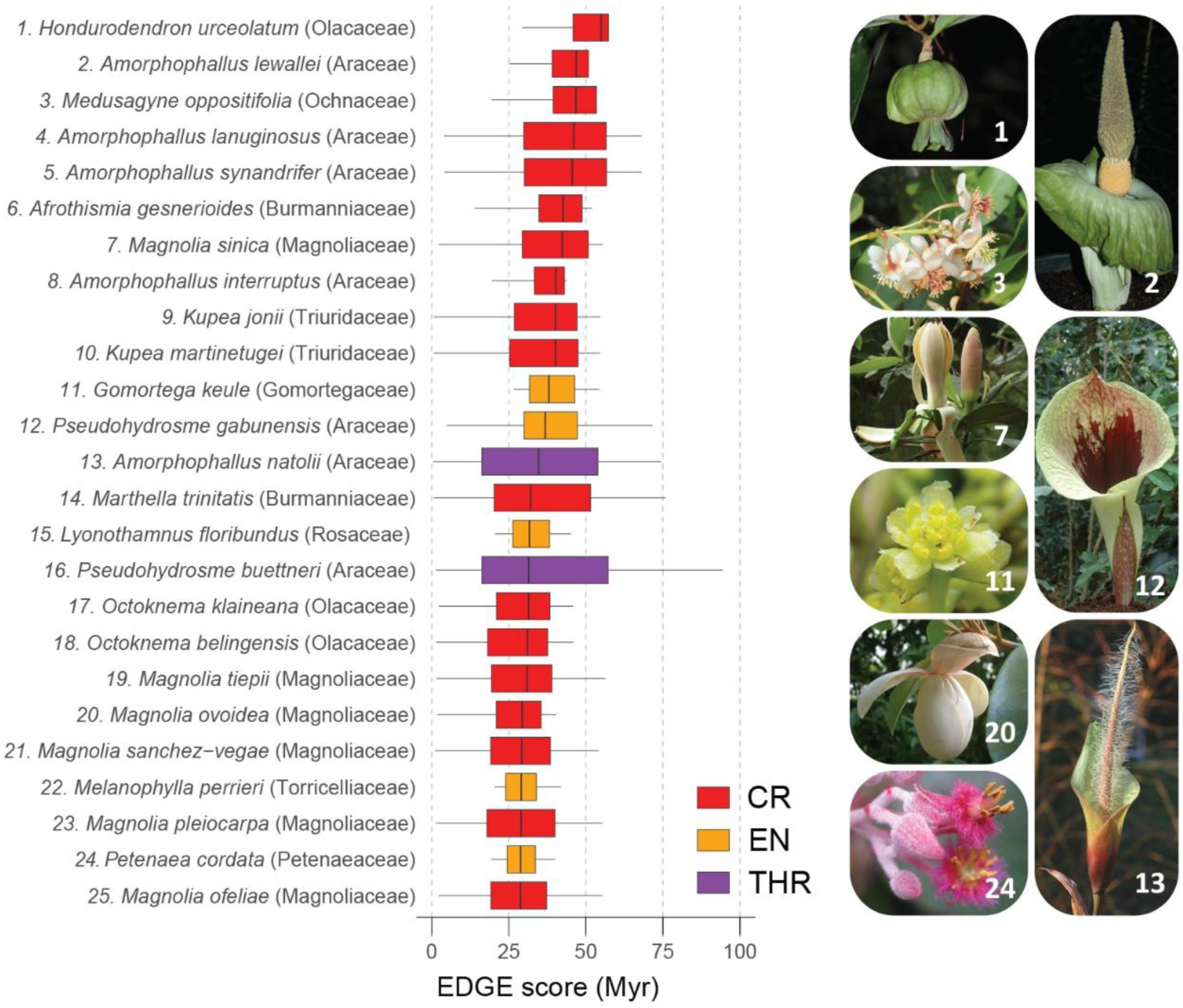
EDGE species. The top 25 Evolutionarily Distinct and Globally Endangered (EDGE) species of angiosperms based on their median EDGE score (boxplot centre line) obtained from the 200 replicates (millions of years, Myr). Boxplots are coloured according to the species’ threat status based on IUCN Red List categories or extinction risk predictions (red, Critically Endangered - CR; orange, Endangered - EN; purple, estimated to be threatened based on extinction risk predictions - THR). Image credits: 1. J. Kolby; 2, 12, 13. W. Hetterscheid; 3, 11. M. Christenhusz; 7, 20. W. Sun; 24. M. (B.) Vorontsova.

The 9,945 EDGE species belong to 259 of the 417 families of angiosperms, with more than 50% of all EDGE species concentrated in 14 families; 31.2% are found within just five: Araceae (893 spp), Myrtaceae (714 spp), Fabaceae (555 spp), Orchidaceae (546 spp) and Annonaceae (393 spp). Unsurprisingly, three of these families also have among the largest amount of evolutionary history at risk, i.e. Araceae, Myrtaceae and Fabaceae (see above). Among families with more than 100 species, the families with the greatest percentage of EDGE species are Magnoliaceae (66.9%), Pandanaceae (37.6%), Burmanniaceae (30.8%), Aquifoliaceae (27.5%) and Xyridaceae (25.3%). A further 38 families have at least 25% of their species on the EDGE list, while Juncaceae is the largest family (473 spp) with no species on the EDGE List.

EDGE species are found primarily in the tropical regions of the world, with Madagascar (950 spp), Borneo (561 spp) and Ecuador (446 spp) the richest botanical countries (Fig. 4). Some of the botanical countries having Mediterranean-type climates also stand out, i.e. Cape Provinces of South Africa (361 spp), Western Australia (224 spp) and Turkey (266 spp; Fig. 4). It is not unexpected that these botanical countries are also where the greatest diversity of EDGE species is found. Given that geographic distribution range size is one of the most important correlates of extinction risk^24,30,44^, and that species endemic to a single botanical country are more likely to be threatened^30^, one would expect that a region harbouring a high diversity of endemic species, as in most of the regions mentioned above^32^, would also have high diversity of EDGE species. In fact, most EDGE species found in these botanical countries are endemic to these regions; 99% of EDGE species found in Madagascar are endemic, 96% in Borneo, Cape Provinces, Western Australia and Turkey, and 82% in Ecuador.

**Figure 4.**
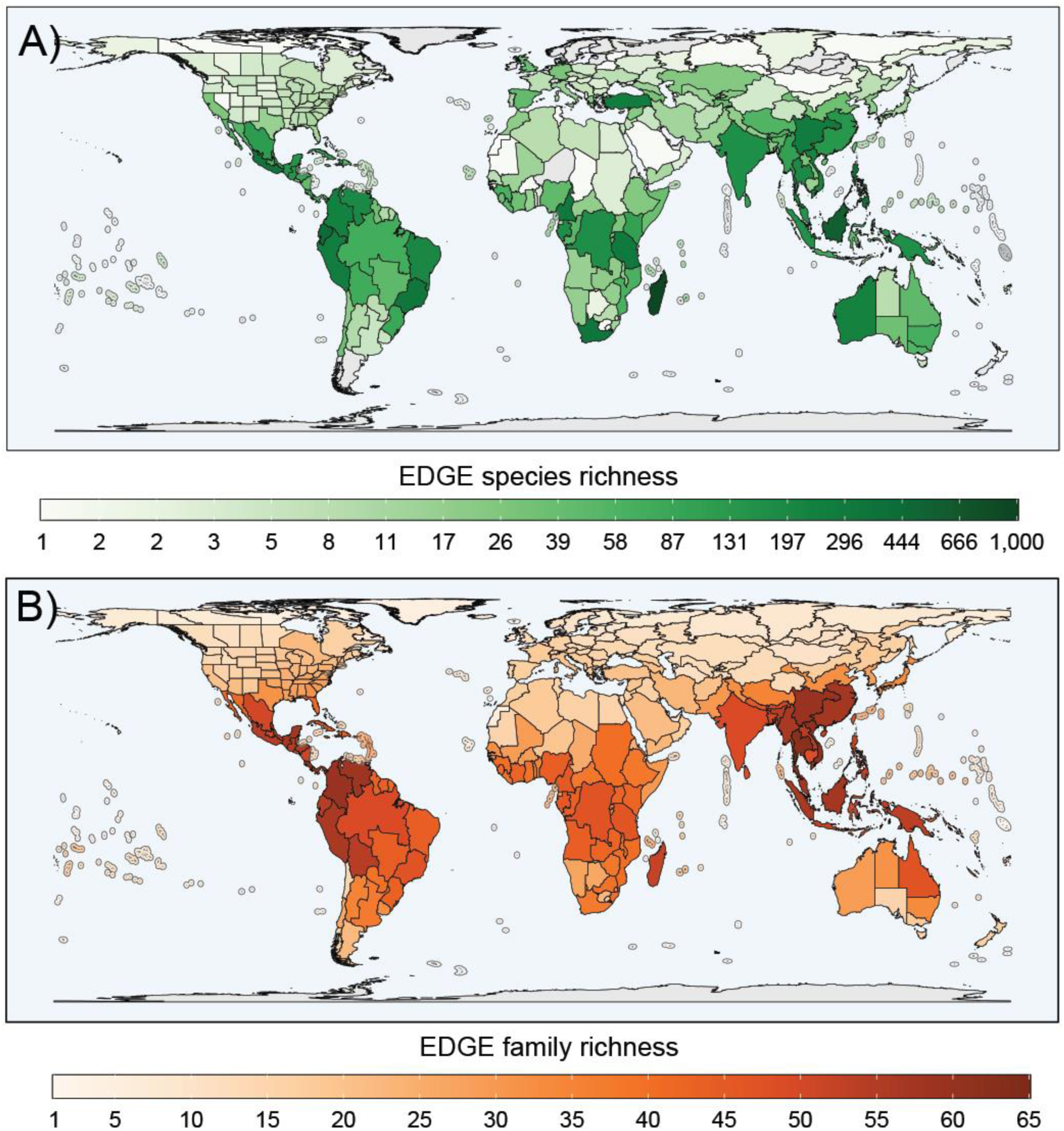
Global distribution of EDGE species and EDGE families. EDGE species richness (A) and EDGE family richness (B) across botanical countries, based on geographical information from the WCVP^28^ and visualised using a Behrmann equal-area projection. Botanical countries represent level three spatial units of the World Geographical Scheme for Recording Plant Distributions^50^.

The EDGE List, obtained through the compilation of EDGE scores using only IUCN Red List assessments (i.e. omitting extinction risk predictions), comprises 4,658 strict EDGE species. Incorporating extinction risk predictions identified a further 1,232 strict EDGE species by accounting for the potential loss of unassessed and data deficient (but likely threatened) close relatives. Of the 4,658 species identified using assessments only, 21 are not on the EDGE list obtained from using both assessments and predictions, a difference of just 0.5%. The high congruence between these two lists demonstrates that the specific value of the extinction risk predictions is twofold: (i) the identification of candidate EDGE species lacking extinction risk assessments and (ii) the addition of strict EDGE species that might otherwise be overlooked due to low assessment coverage of their close relatives.

### EDGE families

The identification of priority groups, termed EDGE lineages, was devised to bring attention to deeper branches of the tree of life that are more at risk of extinction and that may have been neglected by the method focussing on identifying EDGE species^19^. With the large number of EDGE species uncovered in angiosperms, there is also a need to support conservation efforts by being more attentive to families which have a greater proportion of threatened evolutionary history. The concept of EDGE lineage was originally applied at the family level. For a family to be considered an EDGE lineage, it must 1) have a family EDGE score (i.e. mean EDGE score of all species in the family) above the median family EDGE score of all families, 2) have all its IUCN Red List assessed species (except DD) designated as threatened, and 3) have at least half of its species assigned a IUCN Red List category (other than DD)^19^. Given the paucity of IUCN Red List assessments for angiosperms compared with their vertebrate counterparts (e.g. 99.5% of birds vs 20% of angiosperms), we adapted the EDGE lineage approach to identify EDGE families for angiosperms (because only four monotypic families would qualify as EDGE lineages under the original criteria: Cephalotaceae, Eucommiaceae, Gomortegaceae, and Petenaeaceae). An EDGE angiosperm family has (1) a family EDGE score above the median EDGE scores of all families (as in the original EDGE lineage concept^19^), and (2) has a higher-than-median proportion of threatened species, which in the present study is 0.316 (see Methods).

We identify 92 EDGE families using extinction risk predictions (Figs. 2,5; 81 families in the compilation using only IUCN Red List assessments). The most species rich family designated as an EDGE family is Myrtaceae, while the EDGE family with the highest number of EDGE species is Araceae. Seven families identified as EDGE families include no EDGE species, i.e. Brunelliaceae (60 spp), Centroplacaceae (9 spp), Cyrillaceae (11 spp), Cytinaceae (12 spp), Degeneriaceae (2 spp), Dioncophyllaceae (3 spp), and Hanguanaceae (22 spp). These are families that contain large amounts of evolutionary history at risk of being lost from the tree of life despite none of their species being identified as EDGE species. The limited number of species included in the backbone tree in some of these families (e.g. 4/60 spp for Brunelliaceae; 2/11 spp for Cyrillaceae) may increase the variation in EDGE scores among the 200 compilations and result in median EDGE scores below the median (one of the criteria defining EDGE species), which in turn would partly explain why no EDGE species were identified in these families. Increasing the sampling of these groups in the backbone tree in future compilations would be required to assess with more certainty the status of species in these families.

**Figure 5.**
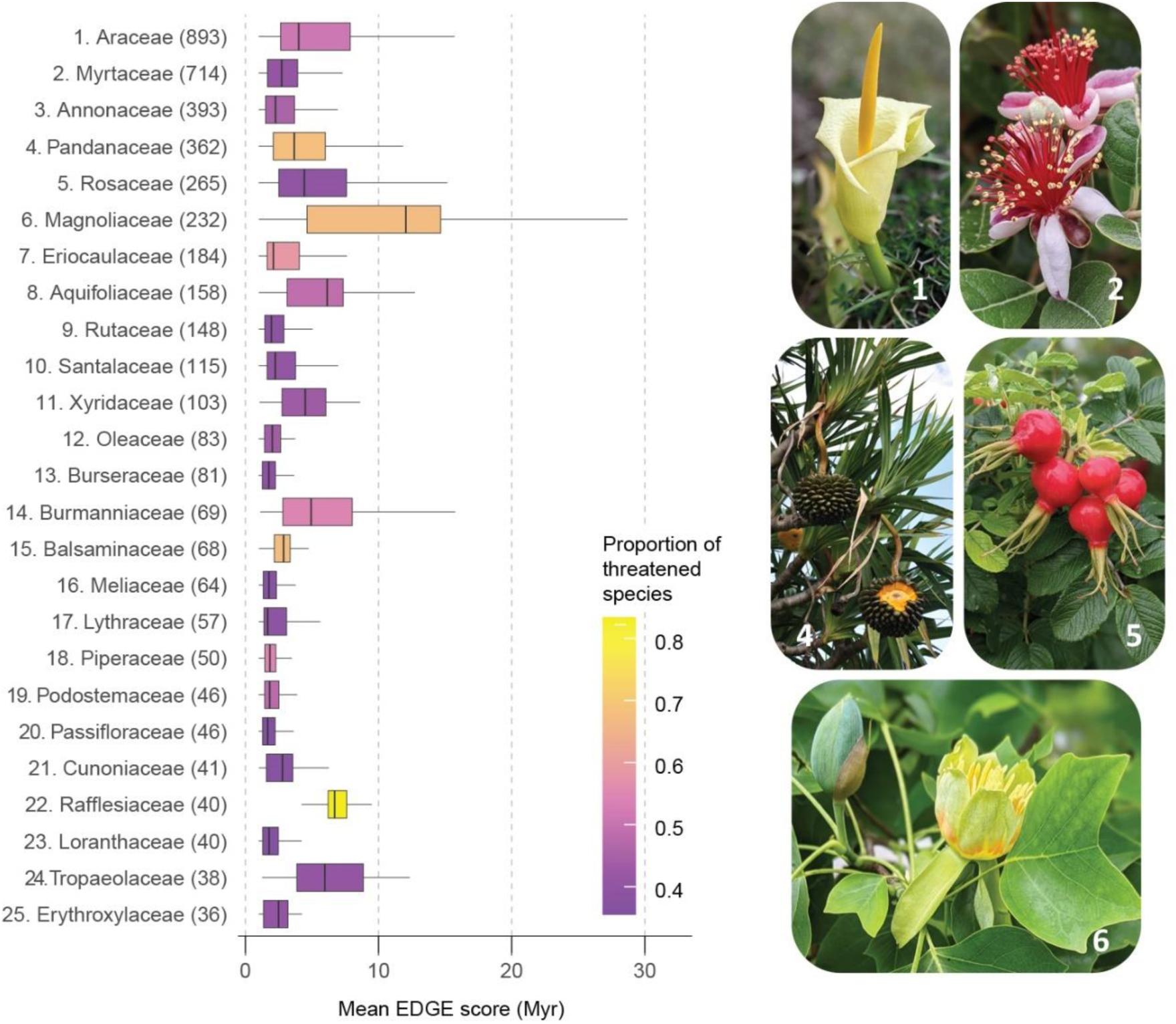
EDGE families. The top 25 EDGE families, ranked by the number of EDGE species they comprise (indicated in bracket). Boxplots show the variation in the mean EDGE score (Myr) of all species in the family across 200 replicates (centre line in boxplots show the median). Boxplots are coloured according to the proportion of threatened species within each family. Image credits: F. Forest.

The distribution of EDGE families is, as for EDGE species, centred in the tropical regions of the world (Fig. 4), with the highest concentrations in South-East Asia, northern South America and Madagascar. The regions with Mediterranean-type climates which are rich in EDGE species are not particularly rich in EDGE families, which suggests that the EDGE species present in these regions are concentrated in relatively few families.

### Threatened useful plants

Plants support most life on Earth, but they also provide resources essential for human well-being. Around one in ten angiosperm species have at least one documented human use, from food and medicines to materials^45^. The preservation of these uses is intimately associated with the conservation of evolutionary history, and this applies also to those benefits that remain to be discovered, referred to as ‘biodiver sity option values^1^’^5,46^. Useful plants represent 20.3% of the total evolutionary history of angiosperms (i.e. 294 byr). Only 5.5% of useful species (i.e. 1,886 species) are estimated to be threatened (vs. 42.4% for all angiosperms based on the present study), the extinction of which would result in the loss of 5.4% of total angiosperm evolutionary history (16.5 byr). While this is a relatively low proportion of useful evolutionary history at risk, it is certainly an underestimate, as it does not account for species whose use is yet to be reported in the scientific literature and documented in digitally available resources. Angiosperms that are evolutionarily distinct are more likely to have a documented human use^13^. However, EDGE species comprise a smaller proportion of useful species than angiosperms more broadly (479 species or 4.8% vs 10.3%), but they represent more than three times (x3.64) as many known useful plants than all threatened species (1,886 species, 1.3%). This higher proportion of useful species among EDGE species compared to threatened species as a whole support findings from previous work showing that conserving evolutionary history is an effective tool for maintaining benefits to humanity^9,11,47^.

### Outlook

Global biodiversity analyses are often dominated by a handful of terrestrial vertebrate groups for which abundant data is available^48^ and the extinction risk of most species has been assessed. The limited availability of such assessments for plants (∼80% of species lack IUCN Red List assessments), exacerbated by gaps and biases in the available data, has led to their exclusion from many global analyses and prioritisation exercises^23^. Here, we used a combination of approaches (e.g. phylogenetic imputation, extinction risk predictions) to address these shortfalls by presenting the first estimate of the global status of threatened evolutionary diversity across the tree of life of flowering plants and identifying the species and families that represent the greatest amount of unique and threatened evolutionary history. Our results directly underpin the two indicators recognised within the Kunming-Montreal Global Biodiversity Framework (GBF)^1^ of the Convention on Biological Diversity, which have been proposed to track progress towards the preservation of evolutionary history, i.e. the Phylogenetic Diversity Indicator and the EDGE index^1,3^.

The potential loss of more than one fifth (i.e. 21.2%) of flowering plant evolutionary history if the current trends are maintained, a proportion much greater than that estimated for vertebrates, illustrates how critical the situation is for plant biodiversity. Failure to slow the current trends of extinction risks, even under the most cautious scenario explored here (i.e. +0.2 category), would result in even greater loss throughout the tree of life of angiosperms (i.e. 28%). Beyond providing the baseline which will eventually allow us to monitor progress in conservation actions through GBF indicators, in particular the EDGE Index, the results reported here will also allow a more targeted conservation approach, with a focus on species and families that are the most evolutionarily unique and most at risk of extinction. For example, the 4,076 candidate EDGE species should be prioritised for formal IUCN Red List assessments. Given that evolutionary history represents the maintenance of options, both for monitoring biodiversity tr ends and to inform sever al aspects of natur e’s contribution to people, it is imperative that plants are included in biodiversity assessments on an equal footing with animals. The present study represents a crucial step towards achieving this goal.

With nearly half of known plant species now estimated to be at risk of extinction^24^, and 75% of those that remain to be discovered predicted to be threatened^49^, how should we prioritise which species are most deserving of immediate conservation action? Our lists of 9,945 EDGE species, and top 19,715 species with highest-ranking EDGE scores that capture 50% of all threatened angiosperm evolutionary history, provide a menu of options for realists who accept that the goal of ‘no species left behind’ may no longer be realistic and that a shorter but mor e compelling list of priorities might be more effective in focusing the efforts of conservation practitioners worldwide.

## Methods

### Species-level phylogenetic trees

We used the GBMB phylogenetic tree of Smith and Brown^51^ as a backbone tree for reconstructing a set of comprehensive species-level trees of angiosperms. This tree comprises 79,874 terminals and is one of the largest phylogenetic trees for seed plants currently available based on DNA sequence data. It has been compiled using the dated phylogenetic tree of Magallon et al.^52^ as backbone and the approach described in Smith and Brown^51^. The names of the terminals in this tree were reconciled to the species list of the World Checklist of Vascular Plants (WCVP; version 11, April 2023^28^). We followed a pragmatic and conservative approach, favouring the removal of a terminal when its taxonomic identity is in any way equivocal. Terminals that could not be assigned to a name recognised by the WCVP were also removed.

After removing all non-angiosperm species (e.g. conifers, ferns) from the tree, we used the function *wcvp_match_names* from the R package rWCVP^53^ to match the remaining 78,927 terminals with accepted names according to the WCVP. This identified terminal names without matches in WCVP, hybrids, unplaced names, subspecific taxa, and duplicates introduced by the assignment of accepted names to non-accepted names, which resulted in 68,171 terminals to be retained in the backbone tree. The other terminals were removed using the function *drop.tip* from the R package ape^54^.

The monophyly of families in the backbone tree was assessed using the R package MonoPhy^55^. Of the 407 families included in the backbone tree, 61 are represented by a single terminal, 332 are found to be monophyletic and 14 are identified as non-monophyletic. The latter were individually examined, and taxa involved in the non-monophyly of families were identified, which resulted in a further 35 terminals removed from the tree. The ten families missing from the cleaned GBMB phylogenetic tree were manually added based on information available in the literature regarding their phylogenetic placement (Table S2). The function *bind.tip* from the R package phytools^56^ was used to include a missing family in the middle of the stem lineage leading to its inferred sister group. The final backbone tree comprised 68,146 terminals.

Missing genera (3,578) and species (267,351) according to the WCVP (version 11^28^) were added using the R package V.Phylomaker^27^ under the scenario 2 option, which assigns missing species to a randomly selected node above the genus-or family-level crown node. Given that we are using the WCVP, which includes families not present in the Plant List (originally used by V.Phylomaker; www.theplantlist.org, v 1.1), a new *nodes.info* file was produced using the backbone tree and the function *build.nodes.1*. The file *nodes.info* contains the genus and family level node information from the input tree required to impute the missing genera and species. The imputation of missing species was repeated 200 times to account for the uncertainty associated with the placement of missing species, which resulted in a set of 200 comprehensive species-level trees of angiosperms with 335,497 terminals.

As mentioned, V.Phylomaker imputes missing species by adding them randomly to nodes within families or genera in the backbone tree. This approach and the large number of missing species creates numerous polytomies, which in turns results in missing species having longer terminal branches on average than species present in the backbone tree and thus inflating their EDGE score. To mitigate this potential bias, we resolve these polytomies using the function *bifurcatr* from the package PDcalc^57^. This approach randomly resolves polytomies, both those found in the backbone tree and those introduced by the imputation of missing species and renders all trees fully binary.

### Extinction risks

We obtained assessments from the IUCN Red List (version 2022-2^37^) for both calculation of EDGE scores and predictions of extinction risk. We reconciled species to accepted species names in WCVP (version 11: April 2023^28^) using *wcvp_match_names* from the R package rWCVP^53^. We used only those assessments based on the current IUCN Red List Categories and Criteria (v3.1^25^) and grouped Data Deficient (DD) species with Not Evaluated (NE). This left us with IUCN Red List categories for 49,800 species, of which 4,380 are Critically Endangered (CR), 8,834 are Endangered (EN), 6,919 are Vulnerable (VU), 3,006 are Near Threatened (NT), 26,552 are Least Concerned (LC), 78 are Extinct (EX), and 31 are Extinct in the Wild (EW).

For the remaining 285,697 species for which data-sufficient IUCN Red List assessments were not available, we predicted their extinction risk status using a Bayesian mixed modelling approach following the methods described by Nic Lughadha et al^30^. We used the WCVP (version 11, April 2023^28^) as a source of life form and distribution data. Species’ native distributions are recorded at the resolution of “botanical countries^5^”^0^, and were filtered to exclude species occurrences coded as introduced, extinct or doubtfully present. We simplified life form data from the modified Raunkiaer system that is used in the WCVP to match the categories used in Humphreys et al^58^. Where species have more than one life form recorded, we assumed that the first life form listed is the most common and used this in our analyses. We scored species as ‘useful’ if they are present in the World Checklist of Useful Plant Species^59^. Following Darrah et al.^60^ and Nic Lughadha et al.^30^, we scored species as ‘threatened’ if they have been assessed as VU, EN, CR, EW or EX by the IUCN Red List^37^ and ‘not threatened’ if they have been assessed as LC or NT. We reconciled species names from these data sources following previously described protocols^30,61^ supplemented in places by manual corrections.

We fitted a Bayesian generalised mixed model to the source data mentioned above using the ‘brms’ package in R^62,63^. We modelled extinction risk as a binary response (threatened or not threatened) drawn from a Bernoulli distribution, with the following formula, where the random effects of ‘endemic_to’, ‘lifefor m’ and ‘continent’ are drawn from the common nor mal distribution:

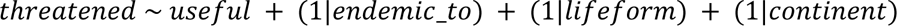

We used weakly informative priors as described in Nic Lughadha et al.^30^; a normal(0,1) distribution for documented human use, half-normal(0,1) distributions for the random effects and a students’ t distribution (3, −1, 1) for the intercept. We sampled 4 chains for 2,000 samples each with 1,000 warmup samples for every model. All models fit well with no divergent samples. We used Pareto-smoothed importance sampling values (PSIS-LOO) and expected log predictive density (ELPD) differences to evaluate model fit, and as in previous iterations^30^, we found that the full model was the best fit (Table S3). We compared these predictions to the observed numbers and proportions of threatened species in each combination of predictors (each^30^) and found that the model generally performs well, but tends to over- and under-predict threat to species at the extremes (Fig. S3) – this is likely due to the limited explanatory capacity of the available predictors. For species-level predictions, we sampled 200 draws from the fitted model, which were used *in toto* to calculate EDGE scores (i.e. species’ predicted extinction r wiasks not averaged prior to EDGE score calculation; see below).

### EDGE score compilation

EDGE scores for each species (i.e. the evolutionary history at risk associated with each species) were compiled using the EDGE2 protocol^5^. The computation of EDGE scores on very large trees, such as angiosperm species-level trees, is computationally challenging, thus the original code has been modified to increase its efficiency (see Code availability). The adapted code restructures the node and edge data of the phylogenetic tree to enable faster data retrieval. Calculations are carried out cumulatively following a tip-to-root traversal, with intermediate results stored within the tree structure. This process avoids repeated calculations and optimizes computational efficiency. The modifications allowed us to compile EDGE scores for all angiosperms, the largest group for which this has been achieved to date.

The probability of extinction for each species was sampled from a distribution of extinction risk values generated by fitting a quartic curve through the median values associated with IUCN Red List categories^5^. Extinct (EX) species were assigned a probability of extinction of 1.0 and Extinct in the Wild (EW) species were assigned the same probability of extinction as CR species. For species without IUCN Red List assessments, we use the extinction risk predicted with the Bayesian mixed modelling approach described above. For species predicted to be threatened, their probability of extinction risk was sampled in the area of the distribution associated with the IUCN Red List categories CR, EN and VU, while for species predicted not to be threatened, it was sampled within the area associated with the LC and NT categories.

To account for the uncertainty associated with the prediction of extinction risk using the Bayesian mixed modelling approach, we assigned a different set of predictions to each of the 200 EDGE score compilations (using a different tree each time) based on a set of 200 draws produced by the Bayesian mixed modelling approach (see above). In each draw, all unassessed species are designated as either threatened (scored 1) or non-threatened (scored 0), while the species with IUCN Red List assessments are assigned their respective categories. This approach allows to account for some of the uncertainty associated with the prediction of extinction risk as each draw represents a statistically plausible set of predictions for all angiosperms.

EDGE species are those that both have an above-median EDGE score (with ≥ 95% confidence) and have been assessed as threatened by the IUCN Red List or by our extinction risk predictions. For the purpose of designating EDGE species in the case of species with extinction risk predictions only (i.e. candidate EDGE species), we use a threshold of 0.5 of draws, i.e. if a species is found to be threatened in 100 or more of the 200 draws, it is considered threatened. Of the 285,697 species without IUCN Red List assessments, 121,977 were predicted to be threatened and 163,720 not threatened based on this threshold. In total, including species with IUCN Red List assessments, 142,219 species are designated as threatened (42.4%) and 193,278 species as not threatened (57.6%).

### Assessing future scenarios

To estimate changes to EDGE scores under scenarios of both subtle and more dramatic increases in global extinction risk, we simulated uniform increases in IUCN Red List category (e.g. all species currently assessed as VU become EN, EN become CR, etc). We incorporated the non-linear increase in probability of extinction associated with each increase in IUCN Red List category^5,6^ by using the curve designed to calculate GE2 scores from IUCN Red List categories (see ‘EDGE score compilation’ above). To avoid stochastic differences that may arise from inter-draw differences, we simulated an increase in GE2 score directly, without resampling from the curve. For each y-value (probability of extinction, sampled per species per draw), we used linear interpolation to project this value to a numeric approximation on the x-axis, where the midpoints of each category are separated by 1 unit (e.g. 0-1=LC; 1-2=NT, 2-3=VU). We then applied a uniform increase to these approximated x-values, where an increase of 1 unit is equivalent to a change of exactly one IUCN Red List category. These adjusted x-values are then reprojected onto the curve, returning an adjusted GE2 score that is equivalent to the same category increase for all species (see Fig. S2). We performed these simulations using an increase of one IUCN Red List category (the most extreme scenario), but this method also allowed us to simulate fractional category increases (0.2 and 0.5 categories), where species are more likely to remain in the same IUCN Red List category, despite a global, uniform increase in extinction probability.

### Quantification of evolutionary history at risk of extinction

We estimate the proportion of the evolutionary history at risk of extinction within angiosperms, which represent the expected loss of phylogenetic diversity, a complementary indicator of the Global Biodiversity Framework^1,3^. We calculated the median expected loss of phylogenetic diversity^64^ for all angiosperms, and the expected loss within each family, using all 200 phylogenetic trees in which each branch is weighted by the probability of extinction of the species it subtends. We compared these values to the median total phylogenetic diversity^15^ of the angiosperms and individual families by summing branch lengths in the original 200 dated phylogenetic trees.

### Useful plants

To determine the amount of threatened evolutionary history represented by plants with at least one human recorded use, we use the list of utilized plant species compiled by Pironon et al^45^, which comprises 35,687 species. Species names from this list of utilized plants were reconciled to accepted species names in the WCVP (version 11, April 2023^28^) using *wcvp_match_names* from the R package rWCVP^53^. This produces an overlapping list containing 34,398 species. We use the same approach as above to compile the proportion of the evolutionary history at risk of extinction within angiosperms represented by useful plant species.

### High-performance computing

The majority of computational analyses outlined here were performed on Crop Diversity HPC at the James Hutton Institute^65^ (https://www.cropdiversity.ac.uk/).

## Acknowledgements

The authors thank Maarten Christenhusz, Wilbert Hetterscheid, Jonathan Kolby, Maria (Bat) Vorontsova, and Weibang Sun for providing images of EDGE species that were included in Fig. 3; Anna Haigh, Isabel Larridon, Jean Linsky, Simon Mayo, Marie-Stéphanie Samain, and Carmen Ulloa for their help in sourcing images; Research Computing at the James Hutton Institute for providing computational resources and technical support for the “UK’s Cr op Diversity Bioinformatics HPC” (BBSRC grants BB/S019669/1 and BB/X019683/1), use of which has contributed to the results reported within this paper. This work benefited from discussions within the IUCN SSC Phylogenetic Diversity Specialist Group (https://www.pdtf.org/).

## Author contributions

FF led the conceptualization of this work; FF, MJMB, SPB, and RG contributed to the methodology; RB, JR, MJMB, FF, RG, and SP contributed to the implementation of computer code; FF, MJMB, and SP performed analyses; FF, MJMB, and RG contribute to data curation; FF, MJMB, and RG wrote the original draft; all authors contributed to the review and editing of the first draft; FF, MJMB, and SP prepared the visualizations; FF was responsible for project administration; SPB, SP, RG, and MJMB contributed equally to this work.

## References

1. Secretariat of the Convention on Biological Diversity. Monitoring framework for the Kunming-Montreal global biodiversity framework. (2022).

2. Díaz, S. et al. Pervasive human-driven decline of life on Earth points to the need for transformative change. Science 366, eaax3100 (2019).

3. Gumbs, R. et al. Indicators to monitor the status of the tree of life. Conserv. Biol. 37, e14138 (2023).

4. Mace, G. M., Gittleman, J. L. & Purvis, A. Preserving the Tree of Life. Science 300, 1707–1709 (2003).

5. Gumbs, R. et al. The EDGE2 protocol: Advancing the prioritisation of Evolutionarily Distinct and Globally Endangered species for practical conservation action. PLOS Biol. 21, e3001991 (2023).

6. Isaac, N. J. B., Turvey, S. T., Collen, B., Waterman, C. & Baillie, J. E. M. Mammals on the EDGE: Conservation Priorities Based on Threat and Phylogeny. PLoS ONE 2, e296 (2007).

7. Altschul, S. F. & Lipman, D. J. Equal animals. Nature 348, 493–494 (1990).

8. Díaz, S. et al. Set ambitious goals for biodiversity and sustainability. Science 370, 411–413 (2020).

9. Forest, F. et al. Preserving the evolutionary potential of floras in biodiversity hotspots. Nature 445, 757–760 (2007).

10. Larsen, F. W., Turner, W. R. & Brooks, T. M. Conserving Critical Sites for Biodiversity Provides Disproportionate Benefits to People. PLoS ONE 7, e36971 (2012).

11. Molina-Venegas, R., Rodríguez, M. Á., Pardo-de-Santayana, M., Ronquillo, C. & Mabberley, D. J. Maximum levels of global phylogenetic diversity efficiently capture plant services for humankind. Nat. Ecol. Evol. 5, 583–588 (2021).

12. Wicke, K., Mooers, A. & Steel, M. Formal Links between Feature Diversity and Phylogenetic Diversity. Syst. Biol. 70, 480–490 (2021).

13. Molina-Venegas, R. Conserving evolutionarily distinct species is critical to safeguard human well-being. Sci. Rep. 11, 24187 (2021).

14. Winter, M., Devictor, V. & Schweiger, O. Phylogenetic diversity and nature conservation: where are we? Trends Ecol. Evol. 28, 199–204 (2013).

15. Faith, D. P. Conservation evaluation and phylogenetic diversity. Biol. Conserv. 61, 1–10 (1992).

16. Gumbs, R., Gray, C. L., Wearn, O. R. & Owen, N. R. Tetrapods on the EDGE: Overcoming data limitations to identify phylogenetic conservation priorities. PLOS ONE 13, e0194680 (2018).

17. Isaac, N. J. B., Redding, D. W., Meredith, H. M. & Safi, K. Phylogenetically-Informed Priorities for Amphibian Conservation. PLoS ONE 7, e43912 (2012).

18. Jetz, W. et al. Global Distribution and Conservation of Evolutionary Distinctness in Birds. Curr. Biol. 24, 919–930 (2014).

19. Gumbs, R. et al. Global conservation status of the jawed vertebrate Tree of Life. Nat. Commun. 15, 1101 (2024).

20. Forest, F. et al. Gymnosperms on the EDGE. Sci. Rep. 8, 6053 (2018).

21. Yessoufou, K., Daru, B. H., Tafirei, R., Elansary, H. O. & Rampedi, I. Integrating biogeography, threat and evolutionary data to explore extinction crisis in the taxonomic group of cycads. Ecol. Evol. 7, 2735–2746 (2017).

22. Pirie, M. D. et al. Spatial decoupling of taxon richness, phylogenetic diversity and threat status in the megagenus Erica (Ericaceae). PhytoKeys 244, 127–150 (2024).

23. Nic Lughadha, Eimear, Antonelli, Alexandre, & Humphreys, Aelys M. Living Planet Report 2020. 36–43 https://www.wwf.org.uk/sites/default/files/2020-09/LPR20_Full_report.pdf (2020).

24. Bachman, S. P., Brown, M. J. M., Leão, T. C. C., Nic Lughadha, E. & Walker, B. E. Extinction risk predictions for the world’s flowering plants to support their cons NerewvaPthioynto. l. 242, 797– 808 (2024).

25. IUCN. IUCN red list categories and criteria: v.3.1. Gland, Switzerland; Cambridge, UK: IUCN. (2012).

26. Ramos-Gutiér rez, I., Lima, H., Vilela, B. & Molina-Venegas, R. A generalized framework to expand incomplete phylogenies using non-molecular phylogenetic information. Glob. Ecol. Biogeogr. 32, 1707–1716 (2023).

27. Jin, Y. & Qian, H. V. PhyloMaker: an R package that can generate very large phylogenies for vascular plants. Ecography 42, 1353–1359 (2019).

28. Govaerts, R., Nic Lughadha, E., Black, N., Turner, R. & Paton, A. The World Checklist of Vascular Plants, a continuously updated resource for exploring global plant diversity. Sci. Data 8, 215 (2021).

29. Rabosky, D. L. et al. An inverse latitudinal gradient in speciation rate for marine fishes. Nature 559, 392–395 (2018).

30. Nic Lughadha, E., et al. Extinction risk and threats to plants and fungi. PLANTS PEOPLE PLANET 2, 389–408 (2020).

31. Alfonzetti, M. et al. Shortfalls in extinction risk assessments for plants. Aust. J. Bot. 68, 466 (2020).

32. Gallagher, R. V. et al. Global shortfalls in threat assessments for endemic flora by country. PLANTS PEOPLE PLANET 5, 885–898 (2023).

33. Borgelt, J., Dorber, M., Høiberg, M. A. & Verones, F. More than half of data deficient species predicted to be threatened by extinction. Commun. Biol. 5, 679 (2022).

34. Butchart, S. H. M. et al. Measuring trends in extinction risk: a review of two decades of development and application of the Red List Index. Philos. Trans. R. Soc. B Biol. Sci. 380, 20230206 (2025).

35. Purvis, A., Agapow, P.-M., Gittleman, J. L. & Mace, G. M. Nonrandom Extinction and the Loss of Evolutionary History. Science 288, 328–330 (2000).

36. Vamosi, J. C. & Wilson, J. R. U. Nonrandom extinction leads to elevated loss of angiosperm evolutionary history. Ecol. Lett. 11, 1047–1053 (2008).

37. IUCN. The IUCN Red List of Threatened Species. Version 2022-2. https://www.iucnredlist.org (2022).

38. Bland, L. M. et al. Cost-effective assessment of extinction risk with limited information. Appl. Ecol. 52, 861–870 (2015).

39. Ulloa, C. U., Nickrent, D. L., Whitefoord, C. & Kelly, D. L. Hondurodendron, a New Monotypic Genus of Aptandraceae from Honduras^1^. Ann. Mo. Bot. Gard. 97, 457–467 (2010).

40. Zuntini, A. R. et al. Phylogenomics and the rise of the angiosperms. Nature 629, 843–850 (2024).

41. Fay, M. F., Swensen, S. M. & Chase, M. W. Taxonomic Affinities of Medusagyne oppositifolia (Medusagynaceae). Kew Bull. 52, 111 (1997).

42. Maas-van De Kamer, H. Afrothismia gesnerioides, another new species of Afrothismia (Burmanniaceae) from tropical Africa. Blumea - Biodivers. Evol. Biogeogr. Plants 48, 475–478 (2003).

43. Muñoz-Concha, D. & Davey, M. R. Gomortega keule, the neglected and endangered Chilean fruit tree. Eur. J. For. Res. 130, 677–693 (2011).

44. Corlett, R. T. The ecology of plant extinctions. Trends Ecol. Evol. S0169534724002805 (2024) doi:10.1016/j.tree.2024.11.007.

45. Pironon, S. et al. The global distribution of plants used by humans. Science 383, 293–297 (2024).

46. Faith, D. P. Valuation and Appreciation of Biodiversity: The “Maintenance of Options” Provided by the Variety of Life. Front. Ecol. Evol. 9, 635670 (2021).

47. Gumbs, R. et al. Conserving avian evolutionary history can effectively safeguard future benefits for people. Sci. Adv. 9, eadh4686 (2023).

48. Ledger, S. E. H. et al. Past, present, and future of the Living Planet Index. Npj Biodivers. 2, 12 (2023).

49. Brown, M. J. M., Bachman, S. P. & Nic Lughadha, E. Three in four undescribed plant species are threatened with extinction. New Phytol. 240, 1340–1344 (2023).

50. Brummitt, R., Francisco, P., Hollis, S. & Brummitt, N. World Geographic Scheme for Recording Plant Distributions. (Hunt Institute for Botanical Documentation, Carnegie Mellon University, Pittsburgh, PA, USA:, 2001).

51. Smith, S. A. & Brown, J. W. Constructing a broadly inclusive seed plant phylogeny. Am. J. Bot. 105, 302–314 (2018).

52. Magallón, S., Gómez-Acevedo, S., Sánchez-Reyes, L. L. & Hernández-Hernández, T. A metacalibrated time-tree documents the early rise of flowering plant phylogenetic diversity. New Phytol. 207, 437–453 (2015).

53. Brown, M. J. M. et al. R WCVP: a companion R package for the World Checklist of Vascular Plants. New Phytol. 240, 1355–1365 (2023).

54. Paradis, E. & Schliep, K. ape 5.0: an environment for modern phylogenetics and evolutionary analyses in R. Bioinformatics 35, 526–528 (2019).

55. Schwer y, O. & O’Meara, B. MonoPhy: a simple R package to find and visualize monophyly issues. PeerJ Comput. Sci. 2, e56 (2016).

56. Revell, L. J. phytools: an R package for phylogenetic comparative biology (and other things): *phytools: R package*. Methods Ecol. Evol. 3, 217–223 (2012).

57. Nipperess, D. An r package for Phylogenetic Diversity (PD) calculations in ecology, biogeography and conservation biology. (2021).

58. Humphreys, A. M., Govaerts, R., Ficinski, S. Z., Nic Lughadha, E. & Vorontsova, M. S. Global dataset shows geography and life form predict modern plant extinction and rediscovery. Nat. Ecol. Evol. 3, 1043–1047 (2019).

59. Diazgranados, M. et al. World checklist of useful plant species. (2020).

60. Darrah, S. E., Bland, L. M., Bachman, S. P., Clubbe, C. P. & Trias-Blasi, A. Using coarse-scale species distribution data to predict extinction risk in plants. Divers. Distrib. 23, 435–447 (2017).

61. Bachman, S. P., Nic Lughadha, E. M. & Rivers, M. C. Quantifying progress toward a conservation assessment for all plants. Conserv. Biol. 32, 516–524 (2018).

62. Bürkner, P.-C. brms: An R Package for Bayesian Multilevel Models Using Stan. J. Stat. Softw. 80, (2017).

63. R Core Team. R: A language and environment for statistical computing. R Foundation for Statistical Computing, Vienna, Austria. (2021).

64. Faith, D. P. Threatened Species and the Potential Loss of Phylogenetic Diversity: Conservation Scenarios Based on Estimated Extinction Probabilities and Phylogenetic Risk Analysis. Conserv. Biol. 22, 1461–1470 (2008).

65. Percival-Alwyn, L et al. UK Crop Diversity-HPC: A collaborative high-performance computing resource approach for sustainable agriculture and biodiversity conservation. PLANTS PEOPLE PLANET ppp3.10607 (2024) doi:10.1002/ppp3.10607.

