## Supplementary Material for "300 billion years of angiosperm evolution at risk of extinction"

### Alternative lists

**Borderline EDGE species.** This group is similar to the EDGE List but includes species with median above a threshold of 80% instead of 95% of the replicates. Given the uncertainty introduced by the imputation of ca 85% of species missing in the backbone tree, it is reasonable to consider further those species which have an EDGE score that fall slightly short of meeting the relatively stringent threshold defining EDGE List species. Many imputed species are likely to have a greater variation of EDGE scores across the 200 trees because of the randomness of the imputation approach, and this would be especially the case for those assigned to genera that are also absent in the backbone tree, or species part of large genera poorly represented in the backbone tree. We identify a further 32,424 species as borderline EDGE species, in addition to the 9,945 species recorded on the EDGE List. Should any further data collection effort be undertaken aiming at refining EDGE score compilation in angiosperms, it should focus on improving the phylogenetic placement of these species as a priority.

**EDGE Research List.** The DD and NE species with EDGE scores above the median EDGE score in at least 95% of the replicates (i.e. at least 190 of the 200 trees) are included on the "EDGE Research List", which essentially catalogue the species that would be on the EDGE List if they were to be assessed and shown to be threatened. The vast majority of angiosperms species have not been assigned a IUCN Red List category (ca 80% of species are DD/NE) and consequently, DD and NE species would be assigned a probability of extinction equivalent to VU species under the EDGE2 protocol. It is thus not surprising that 4,587 species are included in the EDGE Research List when EDGE scores are compiled without considering the extinction risk predictions. There are 5,797 species that are considered DD or NE by the IUCN Red List and that would qualify for the EDGE Research List in the compilation performed with both IUCN Red List and extinction risk predictions. However, species are assigned an extinction risk prediction in this compilation which influence their ED/EDGE scores. Thus, there is essentially no EDGE Research List in this compilation. It is worth noting that the 4,076 candidate EDGE species assigned to the EDGE list are among these 5,797 species. This demonstrate the importance of refining extinction probability predictions, especially in groups where this information is largely unavailable, and highlights which species should be prioritise for a full IUCN Red List assessment.

**EDGE Watch List.** This recognizes the species that fulfil the above median threshold criterion (95%, or 190 of the 200 trees), but which are not threatened (LC, NT or not threatened according to the extinction risk predictions). This list comprises 2,170 species based on the compilation taking into account extinction risk predictions. While these species are not threatened and might not require conservation attention, they do represent a large proportion of unique evolutionary history that we are required to ensure remains safe. A change from not threatened to threatened among these species could lead to a drastic increase in evolutionary history at risk of extinction and number of EDGE species. For example, the safeguard of many borderline EDGE species might be dependent on the protection of deeper branches in the tree of life of angiosperms contributed by species on the EDGE Watch list.

**Table S1. Assessing scenarios of increased extinction risk.** Increases in threatened evolutionary history and number of EDGE species identified resulting from the increase of the probabilities of extinction by +0.2, +0.5 and +1 IUCN Red List category. Total phylogenetic diversity and threatened evolutionary history (median of 200 replicates) are in billions of years.

| | Total phylogenetic diversity | Threatened evolutionary history (% of total) | No EDGE species (% of total) | No EDGE species retain from main analysis (% of main) |
|---|---|---|---|---|
| Main | 1445.3 | 307.0 (21.2%) | 9,945 (3%) | 9,945 |
| + 0.2 | | 347.1 (24.0%) | 11,380 (3.4%) | 9,391 (94.4%) |
| + 0.5 | | 405.1 (28.0%) | 12,358 (3.7%) | 9,459 (95.1%) |
| + 1.0 | | 504.5 (34.9%) | 13,296 (4%) | 9,366 (94.2%) |

**Table S2.** List of ten families not represented in the GBMB phylogenetic tree of Smith and Brown[1] (following homogenisation of names; see Methods) and their inferred position based on the literature.

| **Family** | **Order** | **Genera (no spp)** | **Position** | **References** |
|---|---|---|---|---|
| Apodanthaceae | Cucurbitales | *Apodanthes* (1); *Pilostyles* (11) | Sister to rest of Cucurbitales | 2,3,4 |
| Cytinaceae | Malvales | *Bdallophytum* (4); *Cytinus* (8) | Sister to Mutingiaceae | 2,5,6 |
| Biebersteiniaceae | Sapindales | *Biebersteinia* (4) | Sister to Sapindaceae | 2,7 |
| Cynomoriaceae | Saxifragales | *Cynomorium* (1) | Sister to Crassulaceae, Haloragaceae, Aphanopetalaceae, Tetracarpaeaceae and Penthoraceae | 2,8,9 |
| Triuridaceae | Pandanales | *Kihansia* (2); *Kupea* (2); *Lacandonia* (2); *Peltophyllum* (2); *Sciaphila* (49); *Soridium* (1); *Triuridopsis* (2); *Triuris* (4) | Sister to Stemonaceae, Cyclanthaceae and Pandanaceae | 10 |
| Mitrastemonaceae | Ericales | *Mitrastemon* (2) | Sister to Diapensiaceae | 8 |
| Physenaceae | Caryophyllales | *Physena* (2) | Sister to Asteropeiaceae | 2.8 |
| Rafflesiaceae | Malpighiales | *Rafflesia* (41); *Rhizanthes* (4); *Sapria* (4); | Sister to Euphorbiaceae | 11,12 |
| Setchellanthaceae | Brassicales | *Setchellanthus* (1) | Sister to core Brassicales | 13 |
| Tiganophytaceae | Brassicales | *Tiganophyton* (1) | Sister to Bataceae and Salvadoraceae | 13 |


1. Smith, S. A. & Brown, J. W. Constructing a broadly inclusive seed plant phylogeny. Am J Bot 105, 302–314 (2018).
2. Li, H.-T. et al. Plastid phylogenomic insights into relationships of all flowering plant families. BMC Biol 19, 232 (2021).
3. Filipowicz, N. & Renner, S. S. The worldwide holoparasitic Apodanthaceae confidently placed in the Cucurbitales by nuclear and mitochondrial gene trees. BMC Evol Biol 10, 219 (2010).
4. Bellot, S. & Renner, S. S. Exploring new dating approaches for parasites: The worldwide Apodanthaceae (Cucurbitales) as an example. Molecular Phylogenetics and Evolution 80, 1–10 (2014).
5. Hernández-Gutiérrez, R. & Magallón, S. The timing of Malvales evolution: Incorporating its extensive fossil record to inform about lineage diversification. Molecular Phylogenetics and Evolution 140, 106606 (2019).
6. Nickrent, D. L. Cytinaceae are sister to Muntingiaceae (Malvales). TAXON 56, 1129–1135 (2007).
7. Muellner-Riehl, A. N. et al. Molecular phylogenetics and molecular clock dating of Sapindales based on plastid rbcL , atpB and trnL-trnF DNA sequences. TAXON 65, 1019–1036 (2016).
8. Ramírez-Barahona, S., Sauquet, H. & Magallón, S. The delayed and geographically heterogeneous diversification of flowering plant families. Nat Ecol Evol 4, 1232–1238 (2020).
9. Bellot, S. et al. Assembled Plastid and Mitochondrial Genomes, as well as Nuclear Genes, Place the Parasite Family Cynomoriaceae in the Saxifragales. Genome Biol Evol 8, 2214–2230 (2016).
10. Mennes, C. B., Smets, E. F., Moses, S. N. & Merckx, V. S. F. T. New insights in the long-debated evolutionary history of Triuridaceae (Pandanales). Molecular Phylogenetics and Evolution 69, 994–1004 (2013).
11. Davis, C. C., Latvis, M., Nickrent, D. L., Wurdack, K. J. & Baum, D. A. Floral Gigantism in Rafflesiaceae. Science 315, 1812–1812 (2007).
12. Sun, M. et al. Phylogeny of the Rosidae : A dense taxon sampling analysis. J of Sytematics Evolution 54, 363–391 (2016).
13. Swanepoel, W. et al. From the frying pan: an unusual dwarf shrub from Namibia turns out to be a new brassicalean family. Phytotaxa 439, 171–185 (2020).

**Table S3.** Pareto-smoothed importance sampling values (PSIS-LOO) and expected log predictive density (ELPD) model evaluation of the final model (full) and nested models.

| Model | PSIS-LOO | | ΔELPD | SEΔELPD |
|---|---|---|---|---|
| Full | 40641.1 | ±214.3 | 0 | 0 |
| Continent | 40717.7 | ±214.5 | -38.30 | 8.17 |
| Endemic_to | 41344.6 | ±213.6 | -351.72 | 26.04 |
| Useful | 50955.0 | ±144.1 | -5156.95 | 91.93 |

**Figure S1. EDGE families vs other families across angiosperms.** A) Threatened evolutionary history per species against total evolutionary history per species for EDGE families (in red; R = 0.88, p<0.001) and the other families (in black; R=0.91, p<0.001). Regression lines fitted using a linear model and Pearson correlation coefficient. B) EDGE species richness against total species richness for each family (EDGE families in red, other families in black). Both axes are log scale.

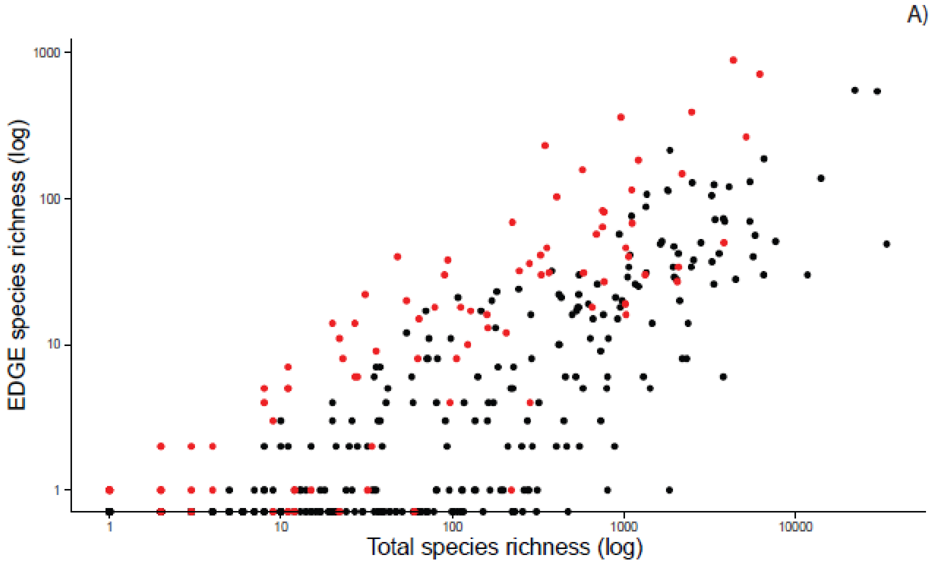


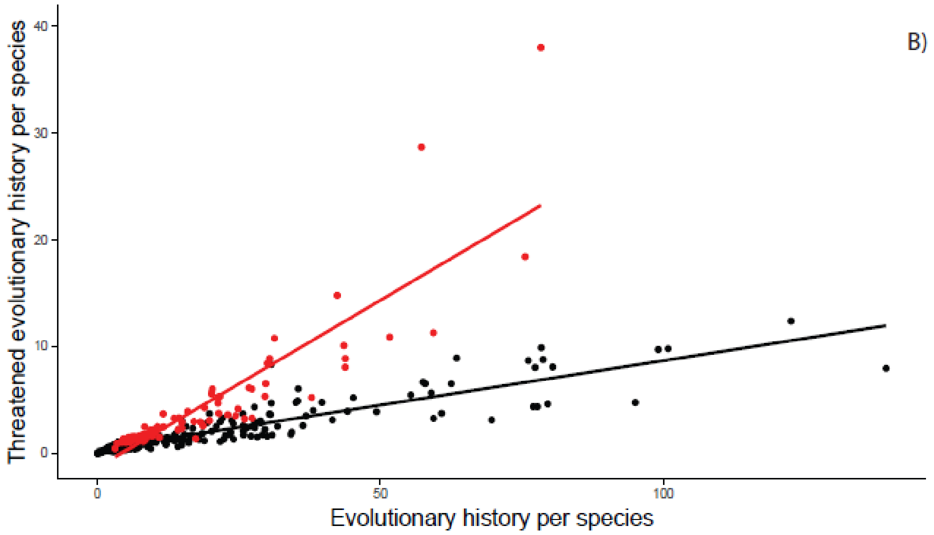

**Figure S2. Simulation of increased extinction risk.** Increase by one IUCN Red List category for two species (only one of the 200 samples shown here, for clarity). Each species is shifted along the x-axis by one category, but the resultant changes in extinction risk are specific to the starting point on the curve. Modified from Fig. 1 of Gumbs et al[5].

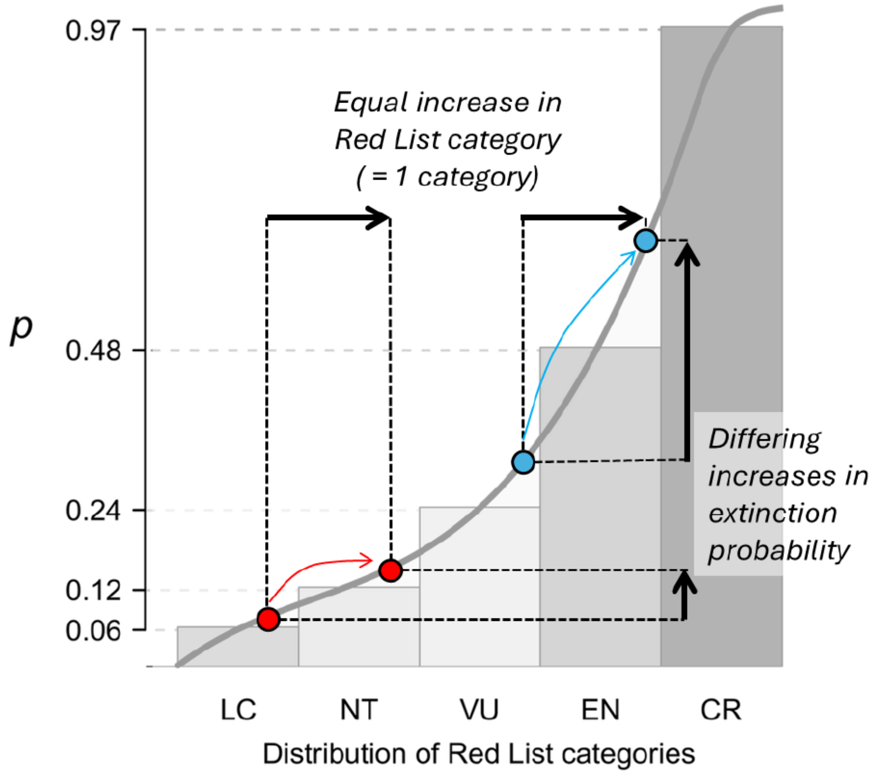

**Figure S3.** Posterior predictive checks to evaluate model fit. Samples from the posterior predictive distribution of the fitted model are used to compare (A) the predicted distribution (blue lines) of values for the proportion of threatened species at each combination of predictor variables to the observed distribution (black line), as well as (B) the prediction distribution (blue lines) of the number of threatened species at each combination of predictor variables to the observed distribution (black line). The samples were also used to compare (C) the predicted number of threatened species at each combination of predictor variables to the observed number.

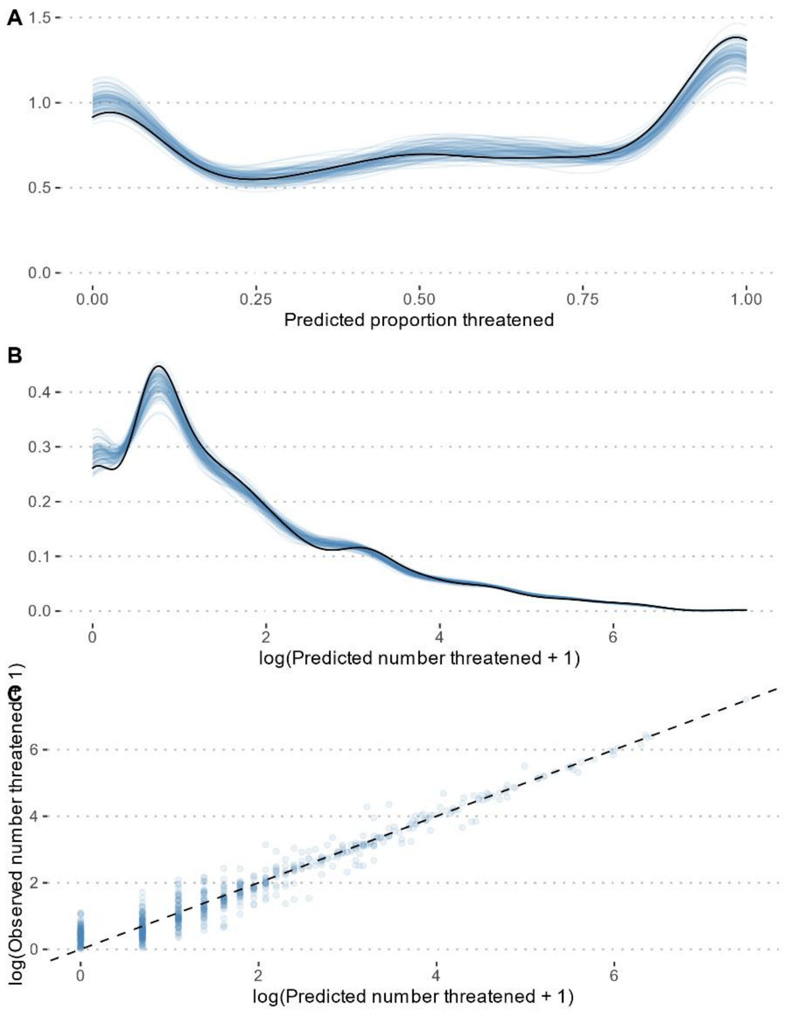
